# CLASS: Accurate and Efficient Splice Variant Annotation from RNA-seq Reads

**DOI:** 10.1101/011718

**Authors:** Li Song, Sarven Sabunciyan, Liliana Florea

## Abstract

Next generation sequencing of cellular RNA is making it possible to characterize genes and alternative splicing in unprecedented detail. However, designing bioinformatics tools to capture splicing variation accurately has proven difficult. Current programs find major isoforms of a gene but miss finer splicing variations, or are sensitive but highly imprecise. We present CLASS, a novel open source tool for accurate genome-guided transcriptome assembly from RNA-seq reads. CLASS employs a splice graph to represent a gene and its splice variants, combined with a linear program to determine an accurate set of exons and efficient splice graph-based transcript selection algorithms. When compared against reference programs, CLASS had the best overall accuracy and could detect up to twice as many splicing events with precision similar to the best reference program. Notably, it was the only tool that produced consistently reliable transcript models for a wide range of applications and sequencing strategies, including very large data sets and ribosomal RNA-depleted samples. Lightweight and multi-threaded, CLASS required <3GB RAM and less than one day to analyze a 350 million read set, and is an excellent choice for transcriptomics studies, from clinical RNA sequencing, to alternative splicing analyses, and to the annotation of new genomes.

## INTRODUCTION

Alternative splicing is an inherent property of eukaryotic genes, with important roles in increasing functional diversity and in disease (1–3). More than 90% of the human genes are alternatively spliced (4,5), with similar levels reported in other eukaryotes. Each gene can produce from one to potentially thousands of splice variants under different cellular conditions, and gene splice isoforms can have similar, independent and even antagonistic functions. Identifying the genes and their splice variations is therefore a critical first step in answering a broad range of biological questions. Over the past five years, next generation sequencing of cellular RNA (RNA-seq) has enabled the discovery of thousands of novel non-coding RNAs and has significantly expanded our catalog of splice variants. However, despite significant progress, extracting gene expression estimates and identifying splice variants in the vast amounts of short read data remains challenging, demanding bioinformatics tools that are fast, accurate and efficient.

The primary goal of a typical RNA-seq analysis is to comprehensively determine the precise exon-intron boundaries on the genome for all transcripts and to estimate their expression levels in the samples. Before this can be accomplished, reads must be mapped to the genome with a fast spliced alignment program that accounts for introns and sequencing errors (reviewed in (6)). Alignments are then pieced together to form gene and transcript models. Virtually all genome-guided transcript assemblers build a graph that represents a gene and its splice variants, and then traverse it to select a subset of transcripts that are likely represented in the sample. Among current programs, Cufflinks (7) connects overlapping reads into overlap graphs, Scripture (8) and IsoLasso (9) build connectivity graphs, and iReckon (10), Scripture and SLIDE (11) generate splice or subexon graphs (reviewed in (12)). Although there are some differences among the sets of parts (exons and introns) predicted by each program, these representations more or less encode equivalent sets of candidate transcripts. Therefore, more important for the program’s accuracy as well as for the number of variants produced is the strategy for selecting transcripts from among the many encoded in the graph. Parsimony-based methods such as Cufflinks’ minimum partition algorithm select a mathematically minimum number of transcripts. They can usually identify the genes and most major isoforms relatively accurately, but are less apt at identifying rarer splicing events. ‘Best fit’ methods, which include IsoLasso, SLIDE and iReckon, choose a subset of transcripts such as to optimize an objective function, using either an integer programming or an expectation maximization formulation. The main problem with these approaches is over-fitting, where programs tend to report a large number of transcripts with very low abundance, most of them spurious. In yet another category, programs such as SpliceGrapher (13) simply omit enumerating transcripts altogether, or otherwise exhaustively enumerate all splice variants encoded in the graph (Scripture). While they can generally capture a larger portion of the true splicing variation, these methods are too imprecise to allow meaningful downstream analyses. Lastly, programs differ in their use of known annotations to inform their predictions. Annotation-guided methods, such as iReckon and SLIDE, rely on an existing set of gene annotations to build their gene models. For species for which there is already an extensive set of gene annotations these methods generally produce more variants, but are also more prone to reporting spurious isoforms and cannot be used to identify novel genes. In contrast, *de novo* programs including Cufflinks, Scripture and IsoCEM, build gene and transcript models from RNA-seq reads alone, without any prior knowledge of gene structure, and therefore are more suited to annotate newly sequenced or less studied organisms. Overall, while many tools already exist to determine the expressed genes and loci in an RNA-seq sample, there is an unmet need for methods that target alternative splicing specifically.

We developed a tool, called CLASS (Constraint-based Local Assembly and Selection of Splice variants), to bridge this gap and detect low abundance splice variation with high accuracy. At its core is the concept of the splice graph, a data structure that we have previously employed in splice variant annotation using both conventional Sanger (EST) (14) and next generation sequencing (15). A splice graph compactly represents a gene with its exons as nodes and introns as edges; splice variants can be read as maximal paths in the graph. CLASS uses a linear programming method to predict exons, and then connects them into splice graphs via introns detected from spliced alignments. Since the splice graph may encode many biologically unfeasible combinations, CLASS uses an efficient dynamic programming optimization algorithm to select candidate transcripts. When compared to reference programs, CLASS captured significantly more splicing variation, both fully reconstructed transcripts and partial splicing events, with high precision. Most importantly, it was the only program tested that produced consistently well formed and easy to interpret annotations for all applications and sequencing strategies. More specifically, our comparative analyses have shown that:

1. CLASS offers the best tradeoff between sensitivity and precision in reconstructing transcripts. In its default setting, it detects 10-70% more transcripts than Cufflinks, which is the most popular and most precise of these programs, at higher or comparable precision; in its sensitive settings, it detects up to twice as many transcripts as Cufflinks for a relatively small drop in precision.
2. It is the best suited to capture local alternative splicing variation. In particular, it can detect up to twice as many alternative splicing events as Cufflinks, with high precision. CLASS finds slightly fewer events than Scripture, which is the most sensitive of the programs, but its precision is considerably (70-80%) higher.
3. It employs a combined gene-level and genome-level model of intronic ‘noise’ that allows more accurate detection of intron retention events.
4. The amount of novel alternative splicing variation detected by CLASS increases with increasingly large data sets.
5. CLASS is multi-threaded and scales well with the amount of data, requiring < 3GB RAM for all of our tests, and can complete a very large run in less than one day.
6. Lastly, since CLASS can produce annotations from RNA-seq data alone, without requiring an existing set of gene annotations, it is very well suited for the annotation of newly sequenced organisms.

We present the overall strategy below, followed by more details about the individual algorithms in the corresponding **Methods** sections and the **Online Supplement**. We then comparatively evaluate CLASS and several popular programs, including both *de novo* and annotation-dependent transcript assemblers, on both control and real RNA-seq sets, in the **Results** section. CLASS is available free of charge for all and under a GNU GPL license from http://sourceforge.net/projects/Splicebox.

## METHODS

### Overview

CLASS determines a set of transcripts in three stages (Figure 1). First, it infers a set of exons from read coverage levels and splice sites using a linear programming technique. Then, it connects the exons into a splice graph via introns extracted from spliced reads. Once the graph is constructed, CLASS selects a subset of transcripts from among those encoded in the graph using an efficient splice graph-based dynamic programming algorithm.

**Figure 1.**
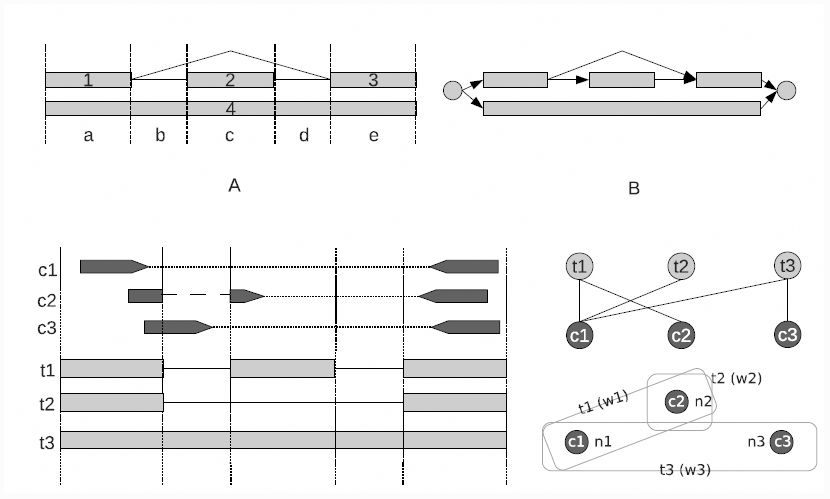
The CLASS transcript assembly algorithm: *Step 1 (A) Exon and introns*. Infer exons from the read coverage levels, using linear programming, and introns from spliced alignments. *Step 2 (B) Splice graph*. Build a splice graph to represent the gene, connecting exons by introns. Shown is a section from a splice graph, with a skipped exon event and a 2-intron retention event, encoding two possible paths (transcripts). *Step 3 (C) Constraints*. Cluster reads into classes (constraints) by their splicing and interval patterns. *Step 4 (D) Transcript selection*. Build and solve the bipartite constraint graph and associated SET_COVER problem. Shown are the constraint graph and SET_COVER problem for four read pairs c1, c2, c3 and c4, and three transcripts t1, t2 and t3.

#### Building the exons

Exons are key to the transcript assembly process, because incorrectly reconstructing exons can miss important gene variations or can create false ones. Since current RNA-seq reads are too short to cover many exons end-to-end, CLASS uses read coverage levels along the genome and splice junctions from spliced read alignments to find exons (Figure 1A and **Methods**). CLASS employs a two-step procedure to determine a set of exons: first, it enumerates all combinations of exons that can explain the splice site patterns and paired-end reads. Second, for each such combination it formulates and solves a linear program expressing several types of constraints. Intuitively, the read coverage levels for all alternative exons over a common interval should cumulatively add up to the observed read coverage levels. Additionally, we assume read coverage levels are locally uniform, and therefore the coverage of adjacent portions of the same exon should be similar. Each exon combination is scored by the linear program, and the combination with the minimum objective function value is chosen in the end.

#### Modeling intronic ‘noise’

Intronic RNA, produced by unspliced transcript that is either residual or part of the experiment, is a common artifact with real RNA-seq samples. Such intronic ‘noise’ can confound the detection of true mRNA resulted from intron retention and alternative transcription start and termination events (16). Distinguishing between ‘signal’ and ‘noise’ is therefore critical for creating a full and accurate set of exons. CLASS models intronic read levels across genomic intervals, both within a gene locus and along the genome, as a statistical Poisson distribution and retains high-scoring intervals (top 2%; p-value = 10e-5) as likely intron retention or alternative gene ends, which are then incorporated into the exon finding procedure.

#### Transcript enumeration and selection

Once a set of exons is determined, CLASS generates a splice graph by connecting the exons (nodes) via introns (edges) extracted from spliced read alignments. Candidate transcripts are encoded in the graph as maximal paths from a node with no incoming edges (source) to a node with no outgoing edges (sink) (Figure 1B). Since the splice graph generally encodes a much larger number of transcripts than is biologically possible, CLASS uses a selection procedure to identify a subset of candidates that can explain all contiguity constraints from spliced reads and paired-reads. In practical terms, a *constraint* is a cluster of reads or read pairs that share the same set of exons or exon fragments and therefore can be assembled into the same transcript (Figure 1C).

Conceptually, we model the problem as a graph with two types of nodes (bipartite), transcripts and constraints, where each transcript node is connected by edges to the constraints it satisfies, and we must select a subset of transcripts that collectively satisfy all constraints (Figure 1D). In early work, we implemented a simple greedy SET_COVER approximation algorithm (15) that aimed to minimize the number of transcripts that could explain all the read patterns, or constraints, without regard to the number of supporting reads. Here we report an improved algorithm that additionally takes into account the read coverage (abundance) information for each transcript and constraint, modeled as a Q(UANTITATIVE)_SET_COVER problem (see **Supplementary Material S1**). It selects a subset of candidate transcripts while simultaneously assigning a set of compatible reads, and it does so efficiently by exploiting the compactness of the splice graph data structure. The algorithm iteratively grows a set of transcripts by selecting, at each step, the transcript that maximizes a scoring function which takes into account both the number of constraints not covered by the current set and their abundance. As a new transcript is selected, reads are simultaneously assigned to it as determined by its set of constraints, and the algorithm is reiterated with the updated sets of constraints and transcripts.

Since the algorithm favors abundant isoforms, transcripts are being selected largely in the order of their abundance, from the most highly to the least expressed. This allows the selection procedure to be terminated whenever the abundance reaches a user-specified cutoff (parameter ‘-F’), with the most trusted isoforms being reported first. In practical terms, it also allows one to run the program in several modes at once. For instance, to more fully capture splicing variation, a user may run the program using a sensitive parameter setting to produce a larger set of potential isoforms, e.g. transcripts whose abundance is at least 1% of that of the most expressed isoform of the gene (‘-F 0.01’). Then, she may filter them with increasing cutoffs to produce more stringent subsets that are usually more precise. Lastly, to further improve the algorithm’s efficiency for genes with complex structures, rather than enumerating all transcripts at each step in the algorithm, CLASS implements an efficient splice-graph based dynamic programming transcript selection procedure, described below. This method considerably reduces both memory and run time, and allows the program to be run on very large data sets without sacrificing sensitivity.

### Exon reconstruction algorithm

To determine the exons, CLASS analyzes regions of the genome covered by reads, which represent exons or combinations of exons, using splice sites to split each region into *intervals*, denoted by letters (a-e) in Figure 1A. Each interval can belong to more than one exon, denoted by numbers, for instance 1-4 in our example. The portion of an exon corresponding to an interval is called *subexon*, for instance 1,a and 4,c. To determine the most likely combination of exons within a region, CLASS enumerates all *feasible* exon sets, i.e. that are necessary and sufficient to explain all splice sites and all reads. For each such set it formulates and solves a linear program (LP), which is used to score the combination. The combination with the best LP score is chosen as the representative set of exons. The linear program is described below.

#### The linear program (LP)-based scoring system for exon combinations

Given a candidate set of exons, CLASS assigns each subexon an (unknown) read coverage level, c_i,j_, defined as the average number of reads per base of subexon *i,j*. Let Cj be the (observed) coverage on interval *j*. We write a linear system with the following constraints: *i) additivity*, meaning that the coverage level in each interval should be roughly equal to the sum of coverage levels of all subexons within that interval, e.g.: | c_1,a_ + c_4,a_ − C_a_ | ≤ ε_a_, *ii) continuity*, i.e. the coverage of subexons of the same exon should be roughly equal, e.g.: | c_4,a_ − c_4,b_ | ≤ ε_4_, *iii*) *conservation*, i.e. the total coverage of all exons should be equal to the total coverage of the region: Σ_i=1,4_ Σ_j=a,e_ C_i,j_ L_j_ = Σ_j=1,e_ C_j_L_j_, where *L_j_* = length of interval j, and *iv*) *non-negativity*: all (sub)exons should be expressed, e.g.: c_1,a_ ≥ 1. The objective is to minimize the total error, Σ *ε*. For single-end reads, this value is used explicitly to score the combination. For paired-end reads, deviations from the observed fragment length distribution are included as penalties to more finely differentiate among likely exon sets. In the end, the exon combination with the smallest score (‘error’) is chosen.

Once determined, exons are connected into a splice graph via introns extracted from spliced alignments, and candidate transcripts are enumerated as maximal paths in this graph. The candidate transcript set is typically much larger than the true set of transcripts. CLASS implements several algorithms to select a subset of transcripts that are the most likely to be represented in the sample, starting from the SET_COVER problem and its variations, as discussed below.

### Transcript selection algorithms

The goal is to select a subset of the transcripts encoded in the splice graph that can collectively explain all the reads, which we formulate as a variation of the SET_COVER problem. We implement an iterative procedure that simultaneously selects the next transcript and assigns reads to it, thus estimating its abundance in the process. To start, we mark the boundaries of the exons along the genomes and divide the gene into intervals, as described above. To reduce space, we group reads (or read pairs) that cover the same set of intervals into classes, called *constraints*. For each constraint *c*_*i*_, we define its *abundance a*_*i*_ as the number of reads (or read pairs) for that constraint divided by the number of possible start positions of the reads within the intervals. Each constraint can be included (satisfied) into one or more candidate transcripts, *c*_*i*_ ~ *t*_*j*_; conversely, a transcript can be viewed as the set of constraints it satisfies: *t*_*j*_ = *{c*_*1*_, …,*c*_*n1*_*}*. We then denote the abundance of a transcript, *A*_*j*_, as the minimum abundance of its set of constraints: *A*_*j*_ = *min {a*_*i*_ | *c_i_~t_j_}*. Let G be a graph with n transcripts *T = {t*_*1*_, …, *t*_*n*_*}* and *m* constraints *C = {c*_*1*_, …, *c*_*m*_*}*. We give a basic enumeration and selection algorithm for relatively simple graphs, and an efficient splice-graph based implementation that can efficiently handle complex graphs below.

#### Basic algorithm

For a small graph, it is feasible to enumerate and assess all candidate transcripts *t*_*1*_, …, *t*_*n*_ encoded in the graph. At each step, the algorithm evaluates all remaining transcripts and selects the new transcript, *t*_*i*_, that maximizes the score function *V*_*i*_ = *n*_*i*_*/(2-A*_*i*_*/max A)*, where ni is the current number of constraints that transcript *t*_*i*_ is compatible with, *A*_*i*_ is the abundance of transcript *t*_*i*_, and *max A* is the maximum abundance over all transcripts of the gene. Once a transcript *t*_*i*_ is selected, its abundance is subtracted from those of the constraints it satisfies: *c*_*j*_ *-= A*_*i*_. If for any constraint the abundance becomes 0, it is removed from the set. The algorithm is reiterated until there are no non-empty constraints.

#### An efficient splice graph-based algorithm

For complex genes that can generate a large number of transcripts, it may not be efficient or even feasible to enumerate and assess all transcripts at each step. Instead, we take advantage of the compactness of the splice graph representation and the locality of the constraints to design a memory and time efficient dynamic programming algorithm. We start by giving an algorithm to iteratively find the next transcript *t*_*i*_ that satisfies the maximum number of constraints *n*_*i*_, by traversing the graph while calculating an optimal path, and then modify it to take into account the abundance, or read numbers.

Let *L* be a subpath (subtranscript) in the splice graph and *L’* the minimum subpath immediately following *L* such that the constraints partially compatible with *L* cannot end after *L’* (‘memory’). We enumerate all the *L’* and recursively calculate the maximum number of constraints *f(L)* of subtranscripts starting with subpath *L*:

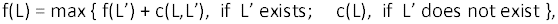

where *c(L,L′)* is the number of constraints partially compatible with *L* (start within *L*) and compatible with the concatenated subpath *L.L′*, and *c(L)* is the number of constraints covered by subtranscript *L*. The algorithm starts with considering every 5’ exon as a subpath. Along with the maximum number of constraints covered, the algorithm can also track the corresponding optimal transcript.

To incorporate abundance information into the optimization process, we modify the algorithm as follows. When processing *L* and *L’*, we exclude subpaths that cover constraints with abundance less than or equal to some fixed value *x*. Hence, the algorithm reports the transcript covering the most constraints among those whose abundance is larger than *x*. We call such a transcript an *x-abundance transcript*. This variation helps determine, at each step, the transcript t* with *max*_*i*_ *V*_*i*_ = *max*_*i*_ *n*_*i*_*/(2-(A*_*i*_*/max A))*. We first calculate the 0-abundance transcript; suppose its abundance is *x*_*1*_. We then calculate the *x*_*1*_-abundance transcript, and so on, until we cannot find any *x*_*m*_-abundance transcript, where x_m_ = *max A*, in the *(m+1)*-st iteration. Then the following Theorem establishes that transcript *t\** is among the transcripts computed.

*Theorem:* The optimal transcript *t\** is among the 0-abundance, *x*_*1*_-abundance, …, *x*_*m-1*_-abundance transcripts.

Proof: Suppose n*, V*, A* corresponds to the number of covered constraints, score and abundance for the optimal transcript t*. Let x_0_=0, then the following two properties hold from the definitions above: (1) 0=x_0_ < x_1_ < … < x_m_; and (2) n_0_ ≥ n_1_ ≥ … ≥ n_m_.

Then A* is between x_0_ and x_m_, and suppose that x_i_ < A* ≤ x_i+1_, where 0≤i<m. Denote the x_i_-abundance transcript by t_i_. Then V_i_ ≤ V*, by virtue of the fact that t* is the optimal transcript. We only need to prove that Vi=V*.

Suppose V_i_<V*, and we already know that n_i_≥n* because the dynamic programming always returns the transcript covering the most constraints (property (2) above). According to the definitions of V_i_ and V*, then it is necessary that A_i_<A* in order to make V_i_<V*. But, A_i_=x_i+1_ based on the definition. Therefore, A*>A_i_=x_i+1_, which contradicts the fact that A* is in the interval (x_i_,x_i+1_]. Hence, the assumption that V_i_<V* is false and we must have V_i_=V* and hence t*=t_i_ (i.e., the x_i+1_-abundance transcript), which concludes the proof.

### Materials and sequences

For our analyses on simulated data, we generated RNA-seq reads with the software FluxSimulator (17), starting from the GENCODE v.17 gene annotations and choosing the options ‘RNA fragmentation’ and 200 million clusters. In total, 15,062 genes and 22,544 GENCODE transcripts were represented by the 200 million 75 bp paired-end reads in the sample. For analyses on real data, strand specific RNA-seq libraries were constructed for the lymphocyte samples using the TruSeq RNA SamplePrep Guide version 15008136_A with modifications. Briefly, for the poly A preparation mRNA was purified from 2 µg of total RNA using Illumina RNA purification beads, the resulting mRNA was fragmented using the Illumina Elute, Prime, Fragment Mix and 1^st^ strand cDNA was synthesized following the TruSeq RNA protocol. Second strand cDNA was synthesized using 8 µl of 10X NEBNext® Second Strand Synthesis (dNTP-Free) Reaction Buffer, 2µl of 10X SuperScript II RT Buffer (NEB), 250uM of each dATP, dUTP, dCTP and dGTP, all of the material from the 1^st^ strand cDNA reaction and 4µl of second strand enzyme (NEB) in a total of 100µl. The reaction was incubated at 16°C for 2.5 hours. The resulting double stranded cDNA was purified, end repaired and adenylated following the TruSeq RNA Sample Prep protocol. One microliter of Illumina adapters were used for the ligation following the TruSeq RNA Sample Prep protocol. The adapter ligated cDNA library was then purified using Ampure beads and subjected to USER enzyme digestion in 5 µl of 10X HotStar PCR buffer (Qiagen), 1 unit of USER enzyme (NEB) in a total of 50µl. This reaction was incubated at 37°C for 15 minutes and the enzyme was inactivated by heating to 95°C for 5 minutes. The digested cDNA was purified using Ampure beads and PCR amplified following the protocol in the TruSeq RNA Sample Prep protocol. For the rRNA-depleted library we started with 5µg of total RNA and removed the rRNA using the Ribo-zero magnetic gold kit (Epicenter) instead of purifying mRNA. Otherwise, the library preparation protocol was identical to the procedure described above. Lastly, for our analyses on very deep sequencing data sets, RNA-seq reads from long RNAs in whole-cell, cytosol and nucleus (2 biological replicates each) of IMR90 lung fibroblast cells were downloaded from the ENCODE project’s web site at UCSC (http://genome.ucsc.edu/ENCODE). All reads were mapped to the human genome hg19 using the software Tophat2 (18) using a combined non-redundant set of GENCODE and RefSeq transcripts as reference annotations and all other default parameters.

### Analysis of alternative splicing events

To evaluate the programs for their ability to capture individual types of alternative splicing events, we generated a reference set of events (exon skipping, intron retention and alternative exon ends) from the simulated data. We used ASprofile (19) to extract events from the transcripts sampled by FluxSimulator, and then filtered them to retain only those actually supported by the reads in the sample. We processed each program’s GTF output in a similar manner and compared against the reference sets. To characterize the sources of errors, we searched the set of false positive predictions from each program against the set of events extracted from the full GENCODE data set, which determine artifacts due to paralogs and splice variants present in the annotation. The remaining false positive events were searched for spurious introns and for class-specific patterns. For intron retention, these include misclassification of 5’ and 3’ terminal exons and of reads from alternative exons overlapping the intron, whereas for alternative exon ends they include spurious chimeric combinations of exon start and ends.

### Evaluation measures

We used conventional measures to assess program performance at the transcript, exon, intron and alternative splicing event levels, including: Recall (sensitivity) = TP/(TP+FN), Precision = TP/(TP+FP), and F-value = 2*Recall*Precision/(Recall+Precision).

## RESULTS

### Comparative evaluation on control data

We evaluated CLASS and several state-of-the-art programs for their ability to reconstruct full transcripts and to capture partial splice variation. We included in our tests four de novo assemblers, namely CLASS (v. 2.0), Cufflinks (v. 2.1.1; (7)), IsoCEM (v. 0.9.1; (9)) and Scripture (v. beta2; (8)), and two annotation-based methods, SLIDE (May 7, 2012 download; (11)) and iReckon (v. 1.0.7; (10)). We ran CLASS in two different modes, stringent (default; ‘-F 0.05’) and sensitive ('-F 0.01'); the latter allows the program to report more minor isoforms. For the annotation-based programs we provided GENCODE v.17 (20) gene annotations as guides. To generate test data, we simulated 200 million 75 bp paired end reads using FluxSimulator (17) and starting from GENCODE v17 gene annotations as models. Reads were then mapped to the human genome hg19 using the program Tophat2 (18) and assembled with each program.

#### Performance of programs in detecting full-length transcripts

To evaluate the performance and also to identify potential limitations and biases of each program we performed two types of analyses. In the first analysis we compared the transcripts produced by each program against the set of transcripts sampled by FluxSimulator, to obtain an unbiased assessment. We then also compared the predictions against a comprehensive set of non-redundant GENCODE and RefSeq transcript models, to identify biases and artifacts due to annotated paralogs and splice variants representing alternative combinations of the same exons. These classes of artifacts are impossible to tease apart on real data, where the ground truth is not known, and will be erroneously counted as true matches, thus over-estimating the program’s performance.

When evaluated against the set of true annotations (Figure 2), most programs detect a majority (63-78%) of the exons and introns (‘set of parts’) of the sampled transcripts, with the notable exception of iReckon, which only finds roughly 52% of the features in each category. SLIDE is the most sensitive among the programs but has very low precision, and Cufflinks and CLASS are the most precise. CLASS and CLASS_F0.01 have the best overall performance, detecting a large fraction of both exons and introns with remarkably high precision, >90% for exons and >97% for introns. Programs rank similarly for reconstructing full-length transcripts. CLASS and CLASS_F0.01 again have the best overall performance as measured by the F-value, a combined measure of sensitivity and precision (see **Methods**), and are able to reconstruct 9% and 16% more full-length transcripts compared to Cufflinks, the next and close runner up.

**Figure 2.**
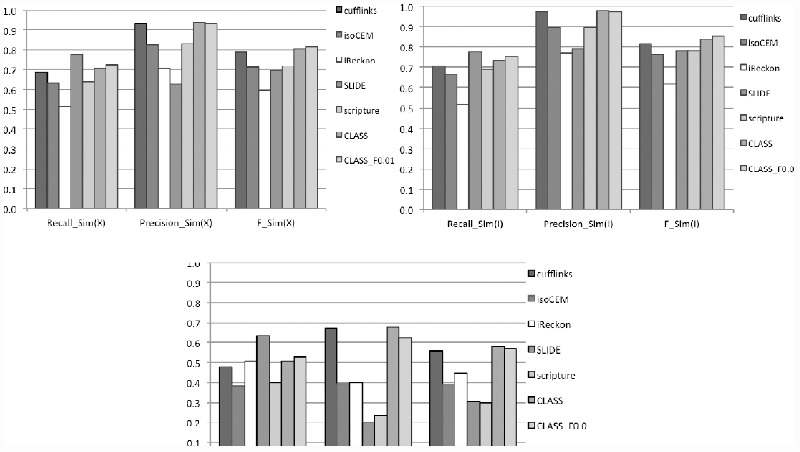
Performance of programs in reconstructing full-length transcripts, on simulated data. Accuracy was measured at exon (X), intron (I) and full transcript (T) levels, by comparison to the subset of reference transcripts sampled by FluxSimulator. Recall = TP/(TP+FN), Precision = TP/(TP+FP) and F = 2*Recall*Precision/(Recall+Precision).

In our second analysis, evaluating the programs against the full set of GENCODE and RefSeq gene annotations revealed several types of biases and errors (Figure 3 and **Supplementary Figure S2**). All programs now seemingly detect the ‘parts’ equally well (~20% sensitivity and 88-100% precision), indicating that many of the false predictions in the earlier comparison come from paralogs of the genes in the sample. Unsurprisingly, programs also seemingly detect more of the reference transcripts, artificially increasing programs’ performance. In particular, the two annotation-based methods show the largest inflation, with SLIDE more than doubling (120% increase) the number of annotation matches and iReckon adding 64% more matches, by virtue of their use of known annotations to scaffold gene models. When we traced these additional matches, most were variants of the sampled genes (53%-92%), and the rest were paralogs (**Supplementary Table S3**), except for iReckon where the variants and paralogs each accounted for roughly half of the false matches. A large portion of the artifacts, between 15% and 67% of the total (with the exception of SLIDE, which had very few), were single exon transcripts. However, even when restricting our analysis to multi-exon transcripts only, SLIDE had very high inflation (128%), followed by iReckon (25%) and Scripture (22%). CLASS (both variations) and Cufflinks had the lowest inflation by far, between 5-7%. Thus, these two programs are the most trusted to produce measurable results on real data.

**Figure 3.**
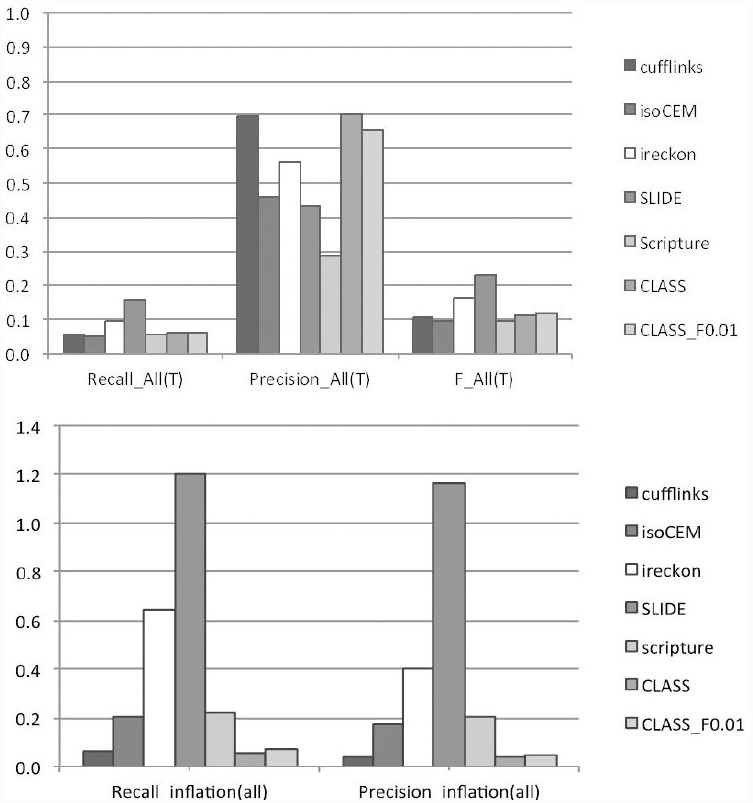
Performance inflation of programs. Observed performance values when measured against the full set of GENCODE reference annotations, for the simulated data (top). Performance ‘inflation’, or the difference between performance measured on the full GENCODE set and the subset of GENCODE transcripts actually represented in the sample (bottom). The additional matches are from spurious paralogs and variants not present in the sample. PCI = *(Match_GENCODE/Match_sim) -1*, where *Match_sim* refers to the subset of transcripts actually present in the simulated sample.

#### Performance of programs in detecting alternative splicing events

Even with the best data, predicting full-length splice variant transcripts from short RNA-seq reads aligned to the genome is prone to assembly errors. Alternative splicing events, which can be determined from the local structure of transcripts or reads, can be detected with more accuracy and are frequently used in studies (21–23). We therefore analyze the ability of the programs to capture primitive classes of alternative splicing events, including exon skipping, intron retention and alternative exon ends. Since most programs do not specifically predict alternative transcription start and termination, we did not include them in the analysis. We compared events detected from transcripts generated by each of the programs to the set of events represented in the simulation data.

As Figure 4 indicates, CLASS_F0.01 and Scripture are the best overall performers as indicated by their F-values, albeit the two programs have strikingly different behavior. Scripture captures the largest number of events in each category, but it does so at the expense of reporting a very large number of false positives, which can severely impact the significance of downstream analyses. CLASS and CLASS_F0.01 find a large portion of the events in each category, balancing sensitivity with high accuracy and achieving the best tradeoff. More specifically, CLASS finds 25-36% more events in each category compared to Cufflinks, which is the leading reference annotation tool and is also the most precise of the programs, at higher or comparable precision. Moreover, CLASS_F0.01 finds roughly twice as many events as Cufflinks in each category with only a relatively small drop in precision (4-17%). Like CLASS, Cufflinks allows users to vary the stringency of the program. We therefore separately compared the performance when varying the parameter range of both CLASS and Cufflinks to control the number of isoforms reported (‘-F f’, with f=0.01, 0.02, 0.03, 0.05, 0.1, 0.1, 0.15). Cufflinks’ performance dropped sharply from its default settings, whereas CLASS showed a consistent performance (**Supplementary Figure S4**). CLASS extended the sensitivity range and, for the same sensitivity level, it delivered significantly higher precision. Therefore, using CLASS in its various settings has the highest potential for applications that involve studies of alternative splicing variation.

**Figure 4.**
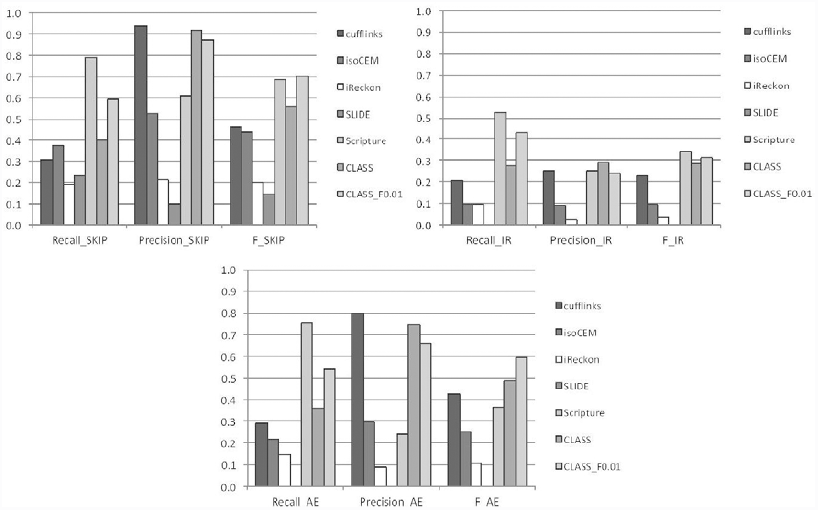
Performance of programs in capturing alternative splicing events: exon skipping (SKIP), intron retention (IR), and alternative exon ends (AE).

We next analyzed the errors made by these programs to evaluate their capacity to capture alternative splicing information. Programs detected exon skipping events with varying degrees of sensitivity (19-79%) and precision (10-94%). Notably, a majority of the false positives for all programs (67-86%; except for SLIDE, 23%) were matches to gene paralogs (Table 1), and only a small fraction were due to other alignment artifacts. This is most clearly illustrated by iReckon and IsoCEM, which predicted large numbers of splicing events, the majority of which were false positives. In contrast, most of the errors for SLIDE were due to spurious introns. The performance of all programs was significantly lower for intron retention events, with 10-52% sensitivity and only 2-29% precision. In most cases, false intron retention predictions resulted from mis-classification of 5’ and 3’ alternative gene starts and ends, as well as from cases in which a splice variant contained an exon that overlapped an intron in the corresponding gene (53-82% of false positives, except for Scripture, 23%). Lastly, programs in general were slightly less accurate in capturing alternative exon ends compared to exon skipping, finding 15-76% of the true variations with 9-80% precision. The errors here were more evenly split between paralogs and variants present in the annotation but not sampled by the data (53-69%; except for iReckon 33%) and from spurious combinations of exon ends. CLASS had both a very low number and a very small percentage of false positives, matched only by Cufflinks, while detecting 30% more features (>90% more when CLASS_F0.01 is used). These analyses also suggest that a simple way in which performance of most programs can be improved is by better distinguishing between true matches and paralogs, and that further improvements can come from better distinguishing between intron retention and other types of variation. Note that the simulated data does not model intronic reads resulted from unprocessed transcripts; the following sections provide a more realistic, albeit empirical, assessment on real data sets.

**Table 1.** Programs’ performance in capturing alternative splicing events. Programs were evaluated for their ability to detect 1,327 exon skipping (SKIP), 185 intron retention (IR) and 1,016 alternative exon end (AE) events present in the simulated data. Incorrect predictions were analyzed to determine classes of artifacts. Artifacts due to paralogs and splice variants of the genes and transcripts in the sample were determined by comparison against events extracted from the full set of GENCODE annotations. The remaining events were searched for spurious introns and for class-specific error patterns, due to mis-classification of alternative first and terminal exons, or of reads from overlapping exons within the same or a different gene (IR), and to spurious combinations of exon start and exon end (AE).

| Program | Predicted | Correct | Recall | Precision | F-value | Artifacts |  |  |
| --- | --- | --- | --- | --- | --- | --- | --- | --- |
| Exon skipping (SKIP) |  |  |  |  |  | Variants+<br>Paralogs | Spurious intron(s) |  |
| CLASS | 586 | 537 | 0.405 | 0.916 | 0.561 | 33 | 6 |  |
| CLASS_F0.01 | 897 | 783 | 0.590 | 0.873 | 0.704 | 92 | 9 |  |
| Cufflinks | 432 | 406 | 0.306 | 0.940 | 0.462 | 20 | 2 |  |
| Cufflinks_F0.01 | 1142 | 782 | 0.589 | 0.685 | 0.634 | 311 | 32 |  |
| IsoCEM | 940 | 496 | 0.374 | 0.528 | 0.438 | 380 | 33 |  |
| Scripture | 1724 | 1045 | 0.787 | 0.606 | 0.685 | 558 | 74 |  |
| iReckon | 1186 | 251 | 0.189 | 0.212 | 0.200 | 781 | 49 |  |
| SLIDE | 3022 | 311 | 0.234 | 0.103 | 0.143 | 618 | 2083 |  |
| Intron retention (IR) |  |  |  |  |  | Variants+<br>Paralogs | Spurious<br>intron(s) | Mis-<br>classified |
| CLASS | 176 | 51 | 0.276 | 0.290 | 0.283 | 17 | 12 | 52+41 |
| CLASS_F0.01 | 331 | 80 | 0.432 | 0.242 | 0.310 | 44 | 57 | 83+63 |
| Cufflinks | 150 | 38 | 0.205 | 0.253 | 0.227 | 19 | 13 | 43+50 |
| Cufflinks_F0.01 | 319 | 68 | 0.368 | 0.213 | 0.270 | 50 | 41 | 89+61 |
| IsoCEM | 205 | 18 | 0.097 | 0.088 | 0.092 | 25 | 61 | 48+51 |
| Scripture | 388 | 97 | 0.524 | 0.250 | 0.339 | 104 | 119 | 49+18 |
| iReckon | 818 | 18 | 0.097 | 0.022 | 0.036 | 116 | 56 | 392+204 |
| SLIDE | 0 | 0 | 0 | 0 | 0 | 0 | 0 | 0 |
| Alternative exon ends (AE) |  |  |  |  |  | Variants+<br>Paralogs | Spurious<br>intron(s) | Spurious<br>combin. |
| CLASS | 496 | 369 | 0.363 | 0.744 | 0.488 | 62 | 4 | 61 |
| CLASS_F0.01 | 831 | 551 | 0.542 | 0.663 | 0.597 | 169 | 11 | 100 |
| Cufflinks | 367 | 293 | 0.288 | 0.798 | 0.424 | 39 | 5 | 30 |
| Cufflinks_F0.01 | 977 | 488 | 0.480 | 0.499 | 0.490 | 326 | 35 | 123 |
| IsoCEM | 761 | 223 | 0.219 | 0.293 | 0.251 | 372 | 40 | 126 |
| Scripture | 3196 | 767 | 0.755 | 0.240 | 0.364 | 1656 | 197 | 576 |
| iReckon | 1721 | 150 | 0.148 | 0.087 | 0.110 | 512 | 50 | 1009 |
| SLIDE | 0 | 0 | 0 | 0 | 0 | 0 | 0 | 0 |

### Comparative evaluation on real data for different sequencing strategies

To assess the performance of programs on real data, we applied them to two large RNA-seq data sets. A lymphocyte sample from an individual free of neuropsychiatric disease was sequenced using two different library preparation strategies, as part of a twin study. In the first method, polyA-selected RNA was sequenced on an Illumina HiSeq2000 instrument to produce roughly 183 million 100 bp paired-end reads. This data set provides a good illustration of a typical RNA-seq analysis experiment, for which most programs are currently optimized. The second library was generated from the same lymphocyte sample by rRNA-depleting the total RNA, and sequenced to generate 317 million paired-end reads. Mapping all reads to the genome with Tophat2 produced roughly 170 million and 240 million read alignments, respectively, but comparatively a larger fraction (46% versus 7%) in the latter sample was in intronic reads.

#### Comparison on the polyA-selected data set

Because the current human genome annotation is inherently incomplete, while also including genes and isoforms not expressed in the sample, it is not possible to determine the true sensitivity and precision of any analysis tool on real data. Nevertheless, we deem consistency with the reference annotation, in particular for sensitivity, as a good indicator of a program’s performance. Using a non-redundant set of GENCODE and RefSeq transcript models as reference, we compare the output of the six programs against the reference annotations. Filtering out single exon assemblies, most of which are biological or computational artifacts, significantly increased the precision of Cufflinks and IsoCEM, whereas there was very little effect on the other programs (**Supplementary Table S5**).

Programs detected between 25-38% of the reference exons and 25-42% of the reference introns, but could only fully reconstruct a small fraction (4-9%; 7,000-16,000) of the annotated transcripts (**Supplementary Figure S6**). This is not unexpected, since only a subset of the reference annotations will be present in any given sample, but the small numbers make it difficult to differentiate among programs and determine the significance. To better assess the relative performance, we designate one method as reference and determine for each of the others the relative change in the number of transcripts found (Figure 5A, top). We chose Cufflinks as reference, because in our earlier testing on simulated data it was the most accurate among the reference programs.

**Figure 5.**
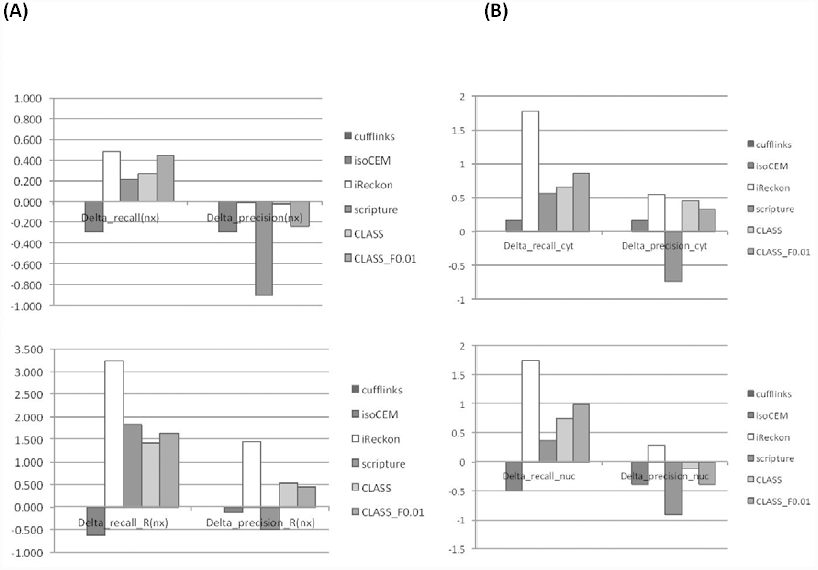
Relative performance of programs on real data. All values are relative to Cufflinks. (A) Performance on two real RNA-seq data sets from lymphocytes from the same individual: a polyA-selected data set (top), and an rRNA-depleted data set (bottom). (B) Performance with very deep sequencing data sets: the ENCODE IMR90 cell line, cytosol sample (top); same cell line, nucleus sample. For a program P, the relative performance improvement for recall is *Delta_recall(P) = [TP(P) – TP(Cufflinks)]/TP(Cufflinks)]*, and similarly for precision. The value for Cufflinks (reference) is 0.

With the exception of isoCEM, programs find 21-48% more transcripts than Cufflinks, with iReckon and CLASS_F0.01 reconstructing the largest numbers of reference transcripts. Cufflinks has the best precision again, followed very closely by iReckon and CLASS. (Note that true ‘precision’ is impossible to assess, as ‘false positives’ could in fact represent true splice isoforms, not found in the reference annotation.) Overall, CLASS and CLASS_F0.01 perform the best among *de novo* assemblers and offer the best tradeoff between sensitivity and precision, as measured by the F-value. When all programs are considered, iReckon appears to perform the best; however, its performance is likely biased by the fact that it used as input the very set of gene annotations we now use for evaluation. When adjusting for paralog and spurious splice variant inflation (**Supplementary Figure S6**, last panel), CLASS and CLASS_F0.01 are the only two programs to exhibit positive cumulative gains in combined sensitivity and ‘precision’ (26% and 41%, respectively, more reference transcripts found compared to Cufflinks, at comparable or slightly lower precision). In conclusion, while Cufflinks appears to be the most precise of the programs for this type of data, CLASS is just as precise while more sensitive, and both CLASS and CLASS_F0.01 offer more accuracy in combined sensitivity and precision.

#### Comparison on the rRNA-depleted data set

We repeated the analysis on the rRNA-depleted RNA sample. Surprisingly, both Cufflinks and IsoCEM performed very poorly, finding only a small subset of reference features; we suspect the reason is that both employ a local intronic ‘noise’ filter at the individual intron level, whereas other programs characterize ‘noise’ at gene (iReckon, CLASS) and/or genome level (Scripture, CLASS). Rankings for other programs were similar to those for the polyA+ data (Figure 5A, bottom; **Supplementary Figure S7**). Although this data set does not fit the characteristics of the simulated data, which was modeled after the polyA-selected RNA sample preparation, we again conjecture that a large portion of iReckon’s performance is in fact due to over-counting of paralogs and alternative exon combinations toward the true matches. CLASS and CLASS_F0.01 are robust with the intronic noise levels and produce reliable gene models, having the best accuracy among *de novo* assemblers. In particular, they can reconstruct 2.5 times as many transcripts as Cufflinks. An example illustrating the programs’ performance at the UBR4-CAPZB locus is shown in **Supplementary Figure S8**.

### Performance of programs on very deep sequencing data sets

The fast and cost effective RNA-seq technology has led to a steady increase in the data size and depth of sequencing, enabling detailed alternative splicing studies. To tackle very large data sets, some programs focus on determining the major isoforms and therefore provide a limited view of the splicing repertoire in a sample, whereas others simply cannot handle the combinatorial explosion. To assess the potential for discovering splicing variation from deep sequencing data sets, we applied all programs to two very large data sets produced by the ENCODE project (24,25). The IMR90 lung fibroblast cell line was sequenced at great depth in three separate surveys, of the whole cell, the cytosolic and the nuclear fractions. Two replicates were run for each fraction, which can be used in our evaluation to assess the accuracy of the predicted features by testing their reproducibility in multiple samples. To reduce the run time, below we restricted our accuracy analyses to chromosome 1. Even so, SLIDE was prohibitively slow and was excluded from the analysis. Summary results of programs are listed separately (**Supplementary Table S9**).

With >300 million reads, the ENCODE IMR90 data sets are among the most deeply sequenced to date and are expected to sample RNA biotypes not found in the reference annotations. Therefore, true accuracy (especially precision) is not possible to assess since novel splice variants will be counted as false positives. Nevertheless, we again judge concordance with annotated features (introns and full transcripts) as indicative of sensitivity and leverage the reproducibility of features across the six samples to better estimate the programs’ performance.

When considering the goal of reconstructing full transcripts, iReckon has seemingly the best performance, as it identified the largest number of transcripts present in the existing annotations (Figure 5B and **Supplementary Figure S10**). Again, however, these results should be considered with caution given the large inflation from variants and paralogs observed with simulated data. Excluding iReckon, both CLASS and CLASS_F0.01 reconstruct the largest number of annotated transcripts in both the cytosol and the nucleus samples, 60-90% (77-103% nucleus) more than Cufflinks and 15-43% (30-49% nucleus) more than the best of the programs, while also having higher or comparable ‘precision’.

We separately evaluated the programs’ accuracy in capturing deeper splicing variation, in particular novel variation, using splice junctions (introns) as surrogates (Table 2). CLASS and CLASS_F0.01 find by far the most known introns, 8% and 11% more than the best of the other programs on the cytosolic sample, and 22% and 37% more on the nucleus sample. When including in the reference those novel introns that are reproducible in at least two data sets, CLASS_F0.01 remained the most sensitive, followed by CLASS and Scripture, at very high precision (>97% for cytosol and >95% for nucleus).

**Table 2.** Performance of programs on the ENCODE IMR90 data: features (full-transcripts and introns) matching known and/or high-confidence novel annotations. GENCODE v17 chromosome 1 annotation contains 15,493 transcripts and 33,202 introns. R = (recall) = Match/Annotations, P = (precision) = Match/Predicted.

| Program | Transcripts |  |  |  | Introns |  |  |  |
| --- | --- | --- | --- | --- | --- | --- | --- | --- |
|  | Predicted | Match | R | P | Predicted | Match | R | P |
| Cytosol |  |  |  |  |  |  |  |  |
| CLASS | 3029 | 1053 | 0.068 | 0.348 | 12662 | 12557<br>(11996) | 0.378<br>(0.361) | 0.968<br>(0.947) |
| CLASS_F0.01 | 3836 | 1183 | 0.076 | 0.308 | 13413 | 13253<br>(12327) | 0.399<br>(0.371) | 0.988<br>(0.919) |
| Cufflinks | 2508 | 621 | 0.040 | 0.248 | 10420 | 10372<br>(10109) | 0.312<br>(0.304) | 0.995<br>(0.970) |
| Cufflinks_F0.01 | 3458 | 719 | 0.046 | 0.208 | 11725 | 11564<br>(10779) | 0.348<br>(0.325) | 0.986<br>(0.919) |
| IsoCEM | 2479 | 722 | 0.047 | 0.291 | 11483 | 11297<br>(10617) | 0.340<br>(0.320) | 0.984<br>(0.925) |
| Scripture | 14621 | 971 | 0.063 | 0.066 | 13820 | 12751<br>(11149) | 0.384<br>(0.336) | 0.923<br>(0.807) |
| iReckon | 4512 | 1730 | 0.112 | 0.383 | 11724 | 11477<br>(10552) | 0.346<br>(0.318) | 0.979<br>(0.900) |
| Nucleus |  |  |  |  |  |  |  |  |
| CLASS | 6084 | 992 | 0.064 | 0.163 | 16391 | 15765<br>(12862) | 0.475<br>(0.418) | 0.962<br>(0.846) |
| CLASS_F0.01 | 10216 | 1141 | 0.074 | 0.112 | 18610 | 17699<br>(14539) | 0.532<br>(0.438) | 0.950<br>(0.781) |
| Cufflinks | 2714 | 561 | 0.036 | 0.207 | 11255 | 11079<br>(10576) | 0.334<br>(0.319) | 0.984<br>(0.940) |
| Cufflinks_F0.01 | 6085 | 789 | 0.051 | 0.130 | 16884 | 16064<br>(13568) | 0.484<br>(0.409) | 0.951<br>(0.804) |
| IsoCEM | 2236 | 277 | 0.018 | 0.124 | 9604 | 8737<br>(7576) | 0.263<br>(0.228) | 0.910<br>(0.789) |
| Scripture | 45247 | 764 | 0.049 | 0.017 | 18048 | 13910<br>(10188) | 0.419<br>(0.307) | 0.771<br>(0.564) |
| iReckon | 5769 | 1539 | 0.099 | 0.267 | 10162 | 9474<br>(8232) | 0.285<br>(0.248) | 0.932<br>(0.810) |

Lastly, CLASS completed the task in roughly 30 min for the chromosome 1 of the cytosol sample and was comparable in speed with the fastest of the programs (**Supplementary Table S11**). As a practical matter, for increased efficiency CLASS_F0.01 can be run first to report a comprehensive set of transcripts, and the output can be filtered using various ‘-F’ parameters (minimum fraction of reported isoforms’ abundance from that of the most expressed isoform) to produce increasingly more precise subsets, at the cost of finding fewer transcripts. Therefore, results for CLASS with multiple settings can be obtained in roughly the same time as a single run.

### *De novo* annotation of a newly sequenced organism

Next generation sequencing has significantly accelerated the pace at which new genomes are being produced. Annotation projects for these genomes are increasingly relying on fast and low cost RNA-seq resources. The choice of RNA-seq transcript assembler here is critical; for instance, since annotation-based programs are not designed to identify novel genes, *de novo* methods are the most productive. To illustrate CLASS’ ability to annotate new genomes, we apply it to enhance the annotation of the peach genome. With its 226.6 MB of sequence assembled in 365 scaffolds, the *Prunus persica* (peach) genome is a good model for future plant species annotation projects. We use CLASS to analyze four RNA-seq data sets sampled from embryo and cotyledon, fruit, root and leaf of peach tree (PRJNA34817), totaling 82.6 MB 75 bp paired-end reads. Preliminary gene annotations are also available, and we use them to identify novel transcript variants that could be used to enhance the existing annotation.

Following read mapping and assembly, CLASS produced between 15,000-27,500 transcript fragments (transfrags) per sample (**Supplementary Table S12**). When compared across the four samples, these amounted to roughly 19,500 transfrags corresponding to existing annotations, but also more than 1000 new loci, each present in at least two of the samples, and 27,161 novel transcripts of known genes, representing new splice variants or extensions of the annotated transcripts (**Supplementary Table S13**). In one example at the *ppa023343m* gene locus (Figure 6A), transfrags assembled from short reads extended the existing gene model by 10-11 exons and revealed several novel splice variations. The extended gene encodes a 1016 aa protein that has similarity over its entire length to importin-11 and importin-11-like proteins in other species (*Prunus mume, Vitis vinifera, Citrus simensis, Fragaria vesca, Theobroma cacao* and *Glycine max*). In another example at the *ppa023750* gene locus, transfrags assembled from the four RNA-seq samples point to additional splice variants, including a novel skipping event of a 39 bp exon located at scaffold_1:4613746-4613784, and a potential retention of an 84 bp intron (scaffold_1:4621242-4621327; Figure 6B), manifested only in the embryo and cotyledon sample. The landscape for this gene is also significantly reconfigured, by merging two previously adjacent genes and by a further extension of its 5’ end. The gene has extensive and close similarity to predicted proteins in apple, Japanese apricot, orange, and cacao. Lastly, a new gene locus, located between genes ppa026188m and ppa005862m, and several putative splice variants discovered with CLASS can be seen in Figure 6C. Blast searches of the two novel putative gene sequences found distant homologs elsewhere in the genome, as well as matches to cytochrome C oxidase subunit 6b protein and to predicted FLX-like proteins in several Rosaceae species. Both sequences contain long open reading frames (762 bp out of 1347 bp, and 234 bp out of the 366 bp sequences, respectively) and are strong candidates for novel, not yet annotated genes.

**Figure 6.**
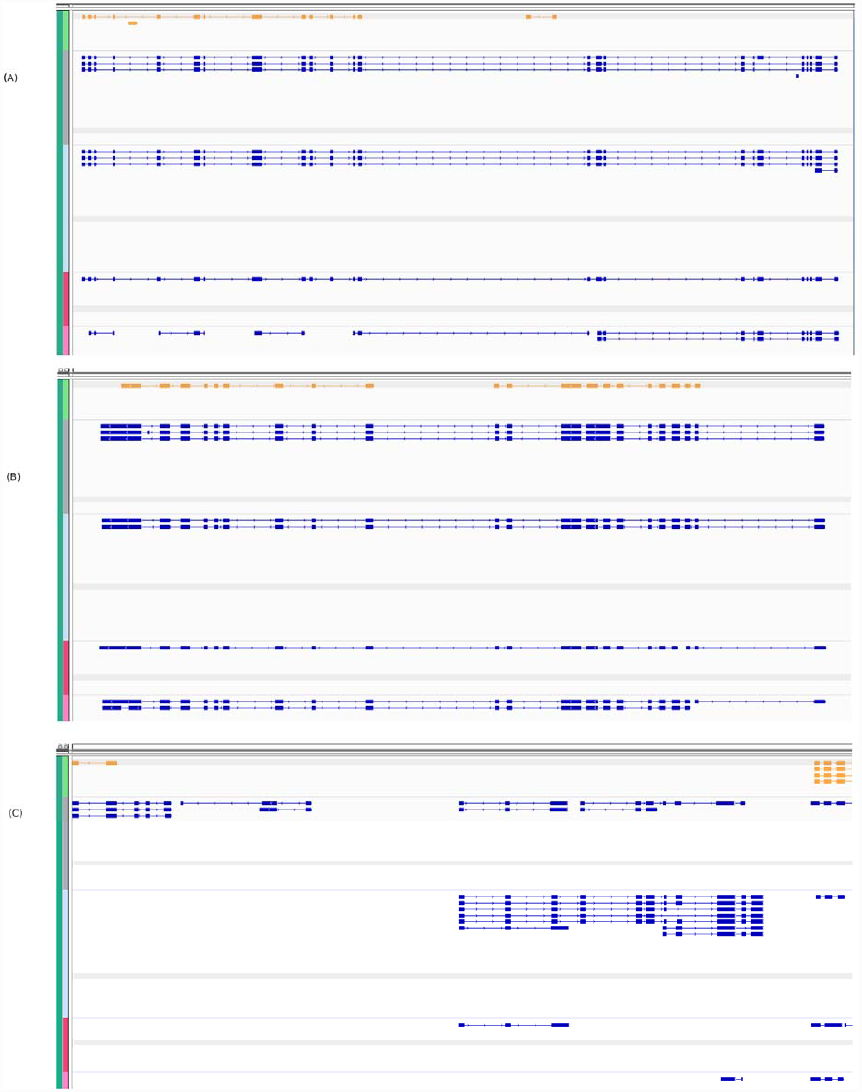
Refining the peach gene models. CLASS transcript predictions for four peach RNA-seq data sets (BioProject ID: PRJNA34817) are shown in blue, and reference annotations in gold. *(A)* RNA-seq reads assembled with CLASS extend the ppa023342m gene model by 10-11 exons and suggest additional splice variants. The extended gene model is supported by data in all of the four samples. *(B)* An extended gene model and several novel splice variants at the ppa023750m gene locus. The intron bridging the two existing gene annotations has (18,7,9,8) supporting reads, respectively, in the four samples, and the last intron is supported by (8,15,6,9) reads. Further, the 39 bp novel exon at Scaffold_1:4613746-4613784 in the SRR531862 sample is alternatively skipped in the reference annotation, and there is ample intronic read support for a putative 84 bp frame-preserving intron retention event at scaffold_1: 4621242-4621327. *(C)* CLASS finds novel genes and splice variants in the intergenic region between annotations ppa026188m and ppa005862m.

## DISCUSSION

A wealth of RNA-seq data, from small individual projects to very large-scale systematic experiments, is making it possible for the first time to catalog alternative splicing variation in detail in different organisms, tissues, at various developmental stages and stress or disease states, and in individual cell types. Many computational methods have already been developed to translate the data into knowledge at the level of genes and transcripts. However, they are still far from being able to assemble full transcript models with high accuracy (26) and have limited ability to capture even local splicing variation, including canonical alternative splicing events. Some classes of events are especially difficult to detect due to artifacts that occur during data generation and mapping (Figure 4 and **Supplementary Table S3**), and have not been systematically pursued by current programs.

We developed a novel splice graph-based algorithm and software tool, CLASS, with the goal to assemble likely models of full-length transcripts while capturing local splicing variations with high accuracy, to allow genome and system-wide alternative splicing analyses. CLASS employs intronic reads and splice junction “noise” models to accurately determine the set of parts, namely exons and introns, and a novel time and memory efficient dynamic programming algorithm to select a subset of probable transcripts that retain most of the splicing variation in the sample.

CLASS differs technically from existing approaches while promoting alternative splicing discovery in several ways: i) it uses an LP-based system to locally predict exon variations, such as alternative 5’ and 3’ exon ends; ii) it incorporates a combined gene- and genome-level model of intronic “noise”, to distinguish retained introns; iii) it models alternative first and last exons, including the cases when they occur at internal exons; and iv) it uses an iterative algorithm and a complex scoring system to select a minimal subset of transcripts that collectively retain as much splicing variation as possible while explaining all the reads.

CLASS also implements several memory and time saving strategies that are critical to its performance and allow it to run on very deep sequencing data sets without sacrificing accuracy. These include a smaller LP system formulated on gene regions rather than along the entire gene, which is both faster and more accurate to solve; clustering reads into classes (‘constraints’); employing a compact and scalable splice graph representation of genes; and, last but not least, implementing a new dynamic programming transcript selection algorithm that avoids enumerating transcripts in complicated graphs, and is memory and space efficient. As a result, a typical run on an Illumina-generated 200 million paired-end read set requires *less than 3 GB* RAM and, when run with multiple threads, takes less than one day, and therefore can be run on most desktop computers.

In our comparative evaluation of CLASS and several state-of-the-art programs, we found CLASS to be significantly more sensitive in capturing alternative splicing variations, both at the level of full transcripts and local alternative splicing events, at precision higher or comparable with that of the best program. In particular, it detected almost twice as much variation as Cufflinks, the most precise of the programs, with only a small decrease in precision. The evaluation also afforded us a unique view of the strengths and limitations of the different approaches. For instance, annotation based approaches as employed by SLIDE and iReckon can detect a larger number of the reference annotations, but are also prone to reporting paralogs and splice variants not actually present in the sample. This is particularly problematic when interpreting the programs’ output on real data, where they would be incorrectly labeled as true matches. The quantity and quality of data can create significant challenges, while library sample preparation can further introduce biases and significantly alter the characteristics of the data (27,28). In general, we found Cufflinks to be the most precise of the programs but missing important splice variations, and Scripture to be the most sensitive but imprecise. However, while different programs may score best by various criteria and for different types of applications, CLASS delivered a consistently good performance for a wide variety of applications and sequencing strategies. These included surveys of polyA-selected (spliced) RNA, which are the most frequent among RNA-seq applications, as well as of ribosomal depleted total RNA, and very deep sequencing experiments to characterize splicing variation, low expression forms, and novel and cellular fraction-specific RNA biospecies, in great depth.

While the boundary between true and noisy splice variation (29) continues to remain undefined, making it ever more difficult to determine the extent of splicing variation and number of isoforms for any given gene, some strategies could help improve the outcome. Better methods are needed to characterize the various types of artifacts that confound classes of variations, such as alternative polyadenylation or alternative promoter usage and retained introns. These can entail implementing sequence models of binding sites of regulatory proteins (30,31), or incorporating other types of evidence including CAGE tags, DNase-seq or FAIRE-seq signals, paired-end diTags (PET-seq) (32) and polyA-seq (33) sequences, where available. Also needed are complete reference data sets on genes or systems that can help evaluate the performance in an unbiased way, or at the very least better simulation models. The latter should include realistic models for sequencing artifacts, including intronic reads from unprocessed pre-mRNA, as well as for the amount and complexity of splicing variation with increasing sequencing depths, and for different types of RNA-seq experiments. Even further, accuracy measures are needed to be able to evaluate programs for their ability to reconstruct splice variations at both global and local levels, including canonical alternative splicing events and local assemblies. Current evaluation schemes focus on the reconstruction of full-transcripts, discounting correct partial reconstructions. Lastly, new sequencing technologies or continuous improvements in the existing ones that extend both read and insert lengths will provide increasing contiguity, while large and judiciously designed experiments will provide multiple replicates or concordant data sets that can be analyzed simultaneously (34,35) to improve both throughput and accuracy.

## FUNDING

This work was supported in part by grants ABI-1356078 and IOS-1339134 from the National Science Foundation to L.F. and a grant from the Stanley Medical Research Institute to S.S..

## ACKNOWLEDGMENTS

We are grateful to Dr. Fuller Torrey for making available the twin samples.

## DISCLOSURE DECLARATION

The authors declare that they have no conflict of interest.

